# ppIRIS: deep learning for proteome-wide prediction of bacterial protein-protein interactions

**DOI:** 10.1101/2025.09.22.677885

**Authors:** Luiz Felipe Piochi, Di Tang, Johan Malmström, Yasaman Karami, Hamed Khakzad

## Abstract

Protein–protein interactions (PPIs) are central to cellular processes and host–pathogen dynamics, yet bacterial interactomes remain poorly mapped, especially for extracellular effectors and cross-species interactions. Experimental approaches provide only partial coverage, while existing computational methods often lack generalizability or are too resource-intensive for proteome-scale application. Here, we introduce ppIRIS (protein–protein Interaction Regression via Iterative Siamese networks), a lightweight deep learning model that integrates evolutionary and structural embeddings to predict PPIs directly from sequence. Trained on curated bacterial datasets, ppIRIS achieves state-of-the-art accuracy across benchmarks while enabling proteome-wide screening in minutes. Applied to Group A Streptococcus (GAS), ppIRIS revealed functional clusters linked to virulence pathways, including nutrient transport, stress response, and metal scavenging. For host–pathogen predictions, ppIRIS recovered 56.2% of known GAS–human plasma interactions with enrichment in complement, coagulation, and protease inhibition pathways. Experimental validation confirmed novel predictions, demonstrating the applicability of ppIRIS for systematic discovery of bacterial and cross-species PPIs. The software is freely available at github.com/lupiochi/ppIRIS.

## Introduction

Protein-protein interactions (PPIs) are fundamental to cellular processes across all domains of life, orchestrating everything from signal transduction and metabolic regulation to immune response and structural integrity [1, 2]. These macromolecular associations form the basis of biological networks that determine cellular function. In host-pathogen relationships, PPIs are particularly critical by mediating processes ranging from adhesion and immune modulation to nutrient acquisition and virulence factor deployment. Understanding these interactions is essential for elucidating disease mechanisms and identifying therapeutic targets

Experimental maps of PPIs such as affinity purification mass spectrometry (AP-MS), chemical crosslinking mass spectrometry, hydrogen-deuterium exchange mass spectrometry, proximity labeling and yeast two-hybrid assays have expanded our understanding of interactomes, but they remain incomplete due to technical limitations [3, 4]. High-throughput methods often struggle with transient interactions, membrane proteins, and proteins expressed at low levels, in addition to false positives and noise. These challenges are magnified for bacterial extracellular effectors and cross-species contacts in physiologic matrices such as plasma, where matrix complexity and throughput constraints limit systematic coverage. Consequently, computational approaches have emerged as valuable complements to experimental techniques.

Recent deep-learning approaches leverage protein language models (pLMs) to infer interaction propensities directly from sequence, improving generalization beyond alignment-based or feature engineering approaches. Sequence-only methods such as D-SCRIPT [5] and TUnA [6] illustrate how pLM embeddings and modern architectures can be effectively utilized for PPI prediction, reporting strong performance on standardized benchmarks. Concurrently, pLMs like ESM-C [7] and ProstT5 3Di [8] provide complementary evolutionary and structure-aware representations at proteome scale. However, most of these predictors have been observed to plateau in performance benchmarks [9], and many are tuned to eukaryotic or model-organism datasets, struggle with bacterial diversity, or are too computationally heavy for rapid proteome-wide and cross-species screening. This creates a need for specialized tools optimized for bacterial contexts, particularly for host-pathogen interaction discovery.

Bacterial pathogens represent one of the leading causes of disease and death worldwide, with over one million deaths associated with drug-resistant infections annually [10]. These pathogens have evolved diverse strategies to invade host tissues, evade the immune system, and establish infections, processes mediated extensively by PPIs. Among these pathogens, *Streptococcus pyogenes*, also known as Group A Streptococcus (GAS), serves as an important case study given its human-specific nature, clinical importance and complex host-pathogen interface.

GAS causes a spectrum of diseases from pharyngitis and impetigo to necrotizing fasciitis and toxic shock syndrome, while also colonizing asymptomatically in many carriers [11]. Historic estimates placed GAS-associated deaths in the hundreds of thousands annually [12], with recent surveillance in the US and Europe indicating renewed surges of invasive disease [13, 14]. As a human-specific pathogen, its transmission, colonization, immune evasion, and dissemination are orchestrated by an extensive virulence repertoire [15]. Many of these virulence factors are secreted or surface-anchored and interact directly with human proteins [16, 17], making GAS an important system for studying host-pathogen PPIs. For GAS and other clinically relevant pathogens, fast and auditable tools that can generalize across strains, score pathogen-host pairs at scale for experimental follow-up are still lacking.

Here, we present ppIRIS (protein-protein **I**nteraction **R**egression via **I**terative **S**iamese networks), a lightweight Siamese model that fuses ESM-C and ProstT5 3Di embeddings to predict bacterial PPIs from sequence alone. Trained on curated bacterial interactions with sequence-identity partitioning, ppIRIS delivers accurate predictions while remaining fast enough for proteome-scale scans. We benchmark ppIRIS against state-of-the-art sequence-based baselines, and then demonstrate its practical utility through application to GAS proteome. In minutes, ppIRIS screens intra-GAS pairs and identifies multiple functional interaction hubs with verifiable structures. We further address the central technical challenge in host-pathogen PPI prediction by introducing an intentional domain shift in the latent space by leveraging different embedding pooling strategies for human and bacterial proteins during training, followed by bridging the two representations for predicting host-pathogen interactions. Through cross-species screening against the human plasma proteome, ppIRIS recovers known biology and nominates high-confidence candidates, with prospective APMS assays supporting a subset of these predictions. Together, these results establish ppIRIS as a practical framework for rapid and auditable PPI discovery in bacterial systems with immediate applicability to GAS and other pathogens.

## Results

### Workflow overview and model architecture

The workflow of ppIRIS starts with comprehensive data collection and preprocessing to ensure high-quality input for the model. We collected PPI data from STRING-DB [18] and bacterial taxonomic information from BacDive [19], focusing on gram-positive bacteria as shown in **Figure 1A**. For model training, we employed an asymmetrical negative-to-positive ratio (5:1 and 10:1), which balances the need for sufficient negative examples while avoiding extreme class imbalance that could impair recall, as described in the methods section. To ensure robust generalization, we implemented a sequence-identity partitioning strategy by clustering bacterial protein sequences with a 40% similarity cutoff to create strictly separated training and validation sets. In addition, we have also utilized the Bernett et al. [20] and D-SCRIPT [5] datasets for benchmarking against state-of-the-art models.

**Figure 1:**
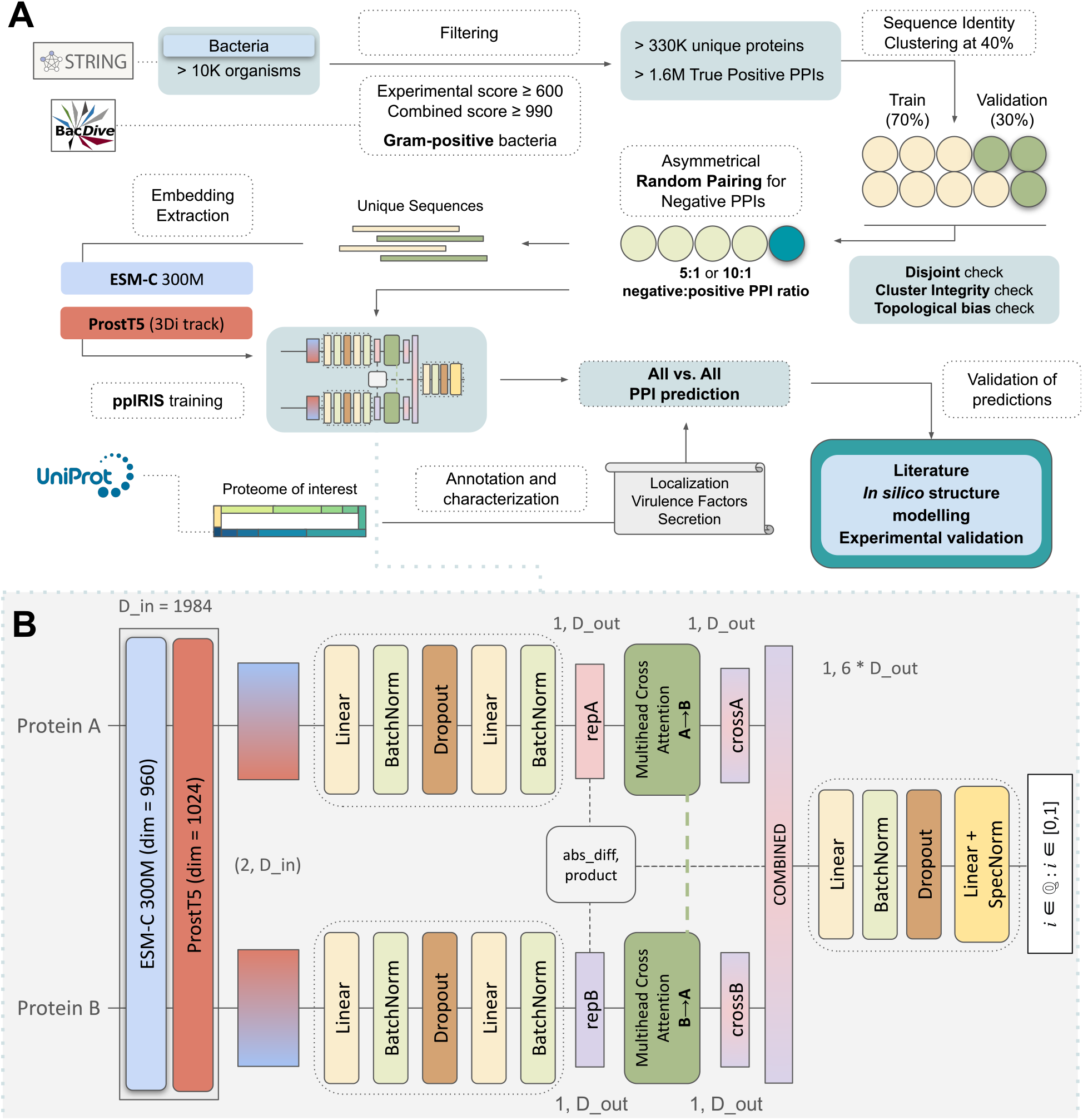
The Workflow and architecture of ppIRIS. **A)** Workflow summary showing data collection from STRING and BacDive databases, protein sequence clustering and negative sampling strategies, and embedding generation with ESM-C and ProstT5 models. In the inference mode, the proteome of interest can be used for the proteome-wide PPI predictions. **B)** The architecture consists of two branches, each processing a different protein sequence from an interaction pair. After computing the input embeddings, each branch passes through a series of fully connected layers, followed by a ReLU activation function. The outputs from both branches are then concatenated and passed through a final fully connected layer to produce the interaction score between 0 and 1.

Depicted in **Figure 1B**, the ppIRIS architecture consists of two main components. First, the Siamese dual encoder approach that leverages complementary protein language model embeddings. Each branch independently processes one protein from an interaction pair through a series of fully connected layers into more compact, information-dense representations while maintaining the unique characteristics of each protein. While previous PPI prediction methods often use a single embedding source, we integrate two state-of-the-art protein language models: ESM-C 300M [7] and ProstT5 [8]. ESM-C provides rich context through masked language modeling on large protein sequence datasets, capturing evolutionary patterns, whereas ProstT5 contributes structural awareness through its bilingual understanding of both sequence and 3D structural features encoded as tokens, particularly through its 3Di track. This combined approach enables ppIRIS to leverage both sequence conservation patterns and structural insights, providing a more comprehensive representation of each protein. The second component of ppIRIS is the feature combination and scoring module. It integrates information from both branches using a fusion strategy that concatenates six distinct feature vectors: the raw encodings from each protein, their cross-attention outputs, their absolute difference, and their element-wise product. This comprehensive approach captures not only the individual protein characteristics but also various relationship patterns between them. The combined representation is then processed through a final network to produce a single interaction probability score. This architecture enables ppIRIS to achieve exceptional computational efficiency compared to existing methods, which is key for proteome-scale predictions, processing millions of potential interactions rapidly while maintaining prediction quality.

### Model performance and benchmarking

To ensure fair and reliable comparisons, we leveraged the comprehensive benchmarking framework established by Reim et al. [9], using a standardized human PPI dataset [20]. We used this dataset to evaluate the performance of both per-residue and per-sequence embeddings (**Supplementary Figure S1**) and showed that per-residue embeddings led to little to no difference compared to per-sequence embeddings for PPI prediction. As shown in **Table 1**, ppIRIS achieved superior performance across multiple metrics, obtaining the highest accuracy (0.661), precision (0.675), and F1 score (0.647) among all compared models. While Richoux-ESM-2 showed slightly better recall (0.654 vs. 0.620), ppIRIS demonstrated a better balance between precision and recall.

**Table 1:** Comparison of model performance metrics across architectures (Reim et al. [9]) using the Bernett et al. human PPI dataset [20].

| Model | Accuracy | Precision | Recall | F1 |
| --- | --- | --- | --- | --- |
| 2d-baseline | 0.575 | 0.630 | 0.373 | 0.468 |
| 2d-Crossattention | 0.641 | 0.660 | 0.589 | 0.623 |
| 2d-Selfattention | 0.616 | 0.611 | 0.553 | 0.591 |
| Richoux-ESM-2 | 0.633 | 0.627 | **0.654** | 0.640 |
| D-SCRIPT-ESM-2 | 0.628 | 0.638 | 0.602 | 0.619 |
| TUnA | 0.645 | 0.672 | 0.580 | 0.622 |
| **ppIRIS** | **0.661** | **0.675** | 0.620 | **0.647** |

Additionally, we also benchmarked ppIRIS using the D-SCRIPT datasets against sequence-based models including D-SCRIPT [5], PIPR [21], Topsy Turvy [22] and TUnA [6], with ppIRIS achieving superior performance across all evaluations (**Figure 2, Supplementary Table S1**). Across all species ppIRIS obtained the highest AUROC and AUPR with gains in AUPR, which is more informative under class imbalance. Relative to the best prior baseline per species, ppIRIS improved on both AUROC and AUPR. Consistent superiority across phylogenetically diverse species suggests that the architecture can capture interaction-relevant invariants rather than overfitting to lineage-specific sequence artifacts.

**Figure 2:**
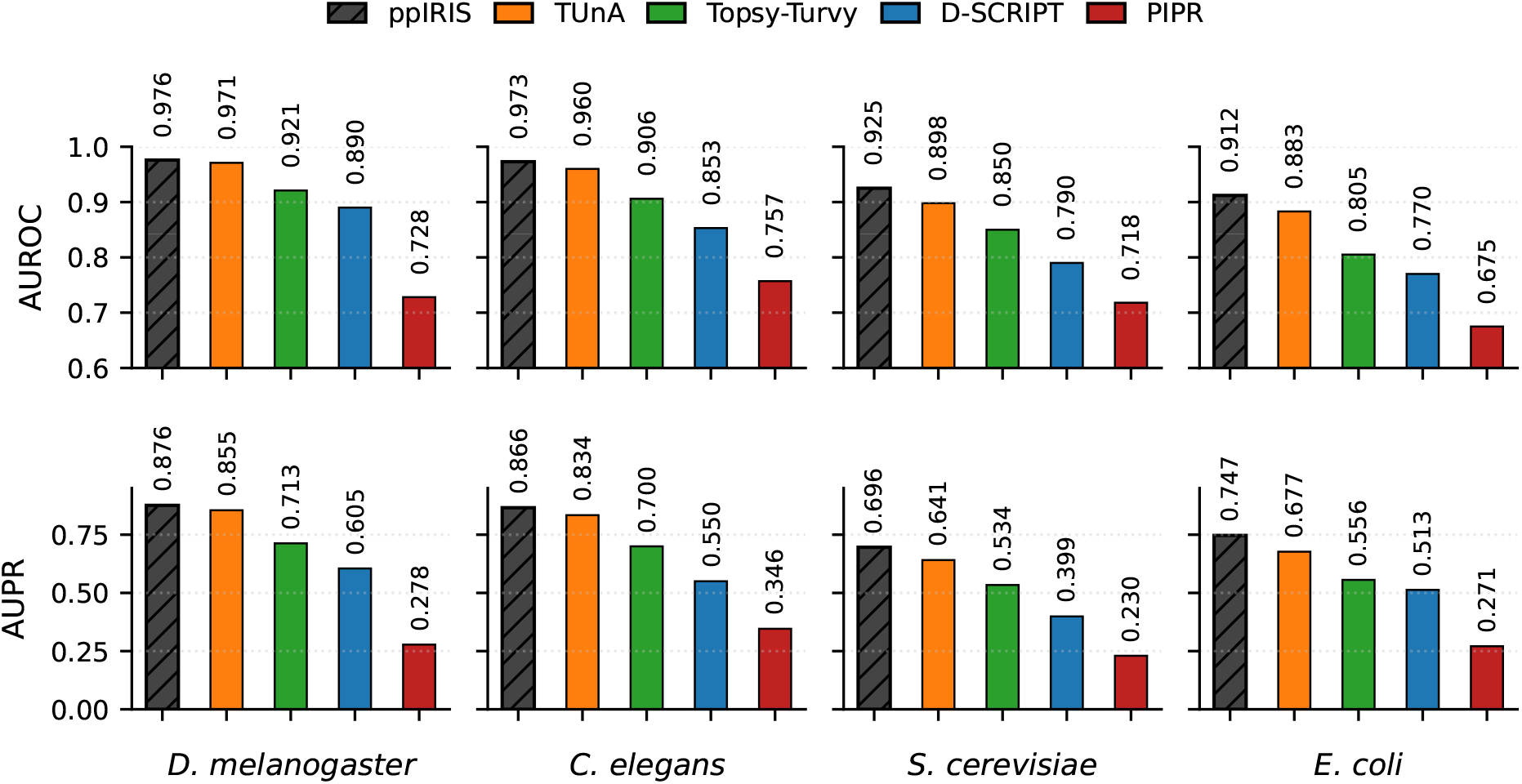
Cross-species AUROC and AUPR comparison on D-SCRIPT benchmark species. Bars show ppIRIS versus prior sequence-only baselines (Topsy-Turvy, PIPR, D-SCRIPT, TUnA). Hatched bar marks the best model per metric. Baseline metrics for species obtained from the original Topsy-Turvy and TUnA.

Having established strong performance on human and mixed-species benchmarks, we turned to gram-positive bacterial PPIs as our primary application domain. Using 40% sequence-similarity partitioning, and combining ESM-C with ProstT5 3Di embeddings yielded the best overall metrics on the 5:1 negative-to-positive split (**Supplementary Table S2** and **Supplementary Figure S2**) (see Methods). ESM-C alone maximized precision, but adding ProstT5 embeddings increased recall and lifted F1 and MCC, indicating complementary signal rather than redundancy. ProstT5 3Di alone underperformed, highlighting sensitivity of purely structure-aware summaries to stronger class imbalance. Across all configurations, the 5:1 ratio split yielded substantially better performance than the more imbalanced 10:1 ratio, with recall being most significantly affected by this imbalance. Ablation analysis showed that replacing the embeddings with random noise matching the embeddings original distribution caused collapse of discrimination (**Supplementary Figure S3**), confirming that performance derives from learned biological representation rather than architectural bias. Together these results show that ppIRIS matches or exceeds heavier baselines on standardized human data and is able to translate across species. In addition, we observed that dual embedding fusion improves bacterial PPI recovery without computational cost of leveraging per-residue pLM embeddings.

### ppIRIS predicts proteome-scale PPIs in minutes

We next applied ppIRIS to the GAS (specifically serotype M1) proteome to explore intra-proteomic interactions. The vast majority of proteins in this proteome have a length of under 500 residues and remain uncharacterized in UniProt (**Supplementary Figure S4**) with no determined structure or annotated function. The model (in inference mode) generated scores for all pairs in less than two minutes (see Methods). As shown in **Figure 3**, our analysis of top-scoring predictions revealed distinct functional patterns when categorized by Gene Ontology (GO) biological process terms. Most known true positive (TP) PPIs for GAS involve ribosomal proteins and metabolic enzymes, which is expected given their well-studied interaction patterns in conserved multiprotein complexes across bacterial species [23]. Thus, we focused on downstream interpretation, by excluding interactions involving ribosomal proteins, re-ranking all pairs by the predicted interaction score and retaining the top 3,000 interactions, which included both known and novel interactions.

**Figure 3:**
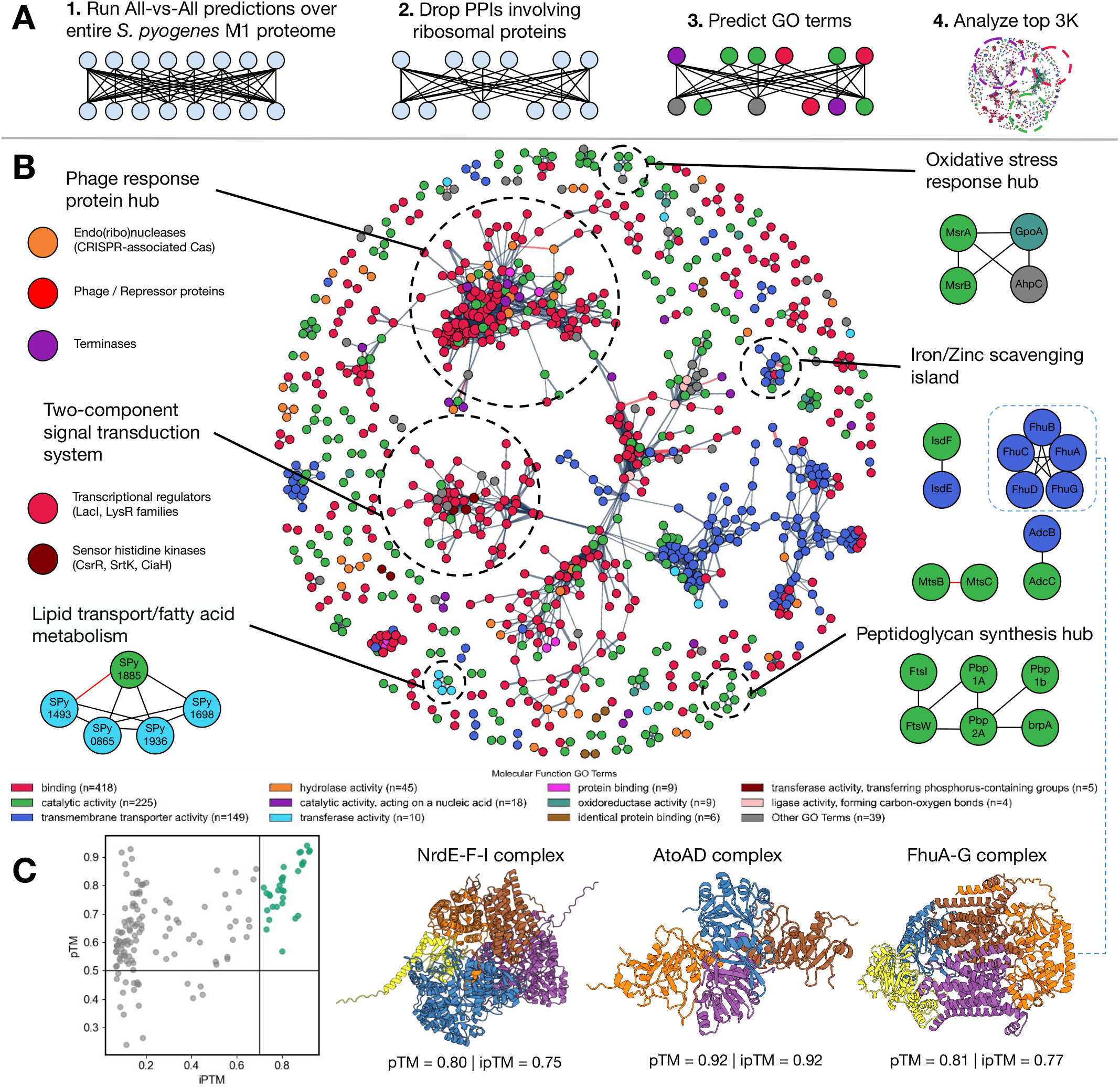
Top intra-proteomic predictions of GAS M1. **A)** Workflow summary: exhaustive scoring, rank-based truncation (top 3,000), ribosomal filtering, functional annotation, and structural validation. **B)** Network of the retained 3,000 highest-scoring non-ribosomal pairs colored by predicted molecular function (DeepGO2); major clusters (transport/metabolism, two-component signaling, phage/defense, metal scavenging) are outlined. **C)** AlphaFold-multimer quality landscape (pTM vs ipTM) for the top 150 modeled pairs; highlighted examples include NrdE-NrdF-NrdI, AtoA-AtoD, and FhuA-FhuG. Green region denotes higher-confidence interface predictions.

Since many GAS proteins remain uncharacterized, we also annotated the GAS M1 proteome using DeepGO2 [24] to predict their molecular function GO terms, which were then used to categorize the predicted interactions. Our predictions predominantly involve proteins annotated with functions related to transmembrane transport, metabolism, stress responses, proteolysis, and gene expression systems (**Supplementary Table S3**). The majority of proteins involved in the interactions have been predicted to have binding, catalytic, and transmembrane transport molecular functions. This highlights ppIRIS’s potential for discovering functional relationships in areas that can be key to bacterial survival and host colonization.

The network visualization in **Figure 3B** also illustrates the complex interaction landscape of the GAS M1 proteome. Different functional clusters were identified and highlighted, such as those involved in lipid transport and fatty acid metabolism, phage response, two-component signal transduction system, and metal scavenging. The phage response and two-component signal transduction system clusters were the largest ones, given the importance of these systems in bacterial survival and adaptation to environmental changes. Other smaller clusters were also identified, such as metal scavenging and peptidoglycan biosynthesis, which are also important for survival in nutrient-limited environments and for maintaining cell wall integrity, respectively. Very few of these interactions have been previously described, making this analysis a valuable resource for further experimental validation and functional characterization.

For structural plausibility assessment, we selected the top 150 scored pairs (with ribosome filtering) and modeled them with AlphaFold-multimer [25]. The predicted complexes involving the NrdE-F-I, AtoAD and FhuA-G protein complexes received the highest pTM and ipTM scores. The NrdE-F-I complex are components of a ribonucleotide reductase in GAS [26] which is essential for DNA synthesis and repair, whereas the AtoAD complex is involved in fatty acid metabolism and has been described in other bacteria [27]. The FhuA-G complex is involved in iron uptake, which is crucial for bacterial survival in nutrient-limited environments, has been characterized in *S. aureus* [28]. Despite these successful cases however, many of the predicted complexes received low pTM and ipTM scores. This can potentially stem from frequent intrinsically disordered regions in bacterial proteins, as well as sparse homology coverage consistent with known AlphaFold-multimer limitations [29].

### Cross-species predictions between GAS M1 and human plasma proteins

We next investigated whether ppIRIS, trained only on intra-species (human-human and bacteria-bacteria PPIs) could generalize to host-pathogen pairs. We combined the gram-positive bacterial PPI dataset with the human PPI dataset from [20] and retrained the model. Using the same dataset creation approach as before (see Methods), we evaluated model performance on a curated GAS-human plasma interactome panel. The evaluation set derived from our previous study [17], comprised 16 well-characterized GAS virulence factors and 62 abundant human plasma proteins (992 possible pairs), of which 112 have prior AP-MS support. As detailed in Methods, we created an intentional pooling-based domain shift during training by using the ProstT5 *< AA*2*F old >* token pooling for bacterial proteins and mean pooling for human proteins, while keeping the same attention-pooled ESM-C representations (**Supplementary Figure S5**). This exposes the model to complementary ProstT5 3Di summaries and encourages reliance on interaction-stable features rather than species-specific pooling artifacts. At inference, we unified the representation by applying a single pooling scheme to all proteins, removing the artificial discrepancy and enabling consistent cross-species scoring (Figure 4A). This representation shift strategy appears to have encouraged reliance on interaction-stable features, as despite no explicit cross-species labels, high-scoring pairs aggregated in biologically relevant pathways, and species-marking artifacts did not dominate clustering (Figure 4A). At a score threshold of 0.5, ppIRIS recovered 63 of the 112 reference interactions (56.2%). Score distributions were shifted for known positives, indicating enrichment rather than random high scoring. The network view (**Figure 4B**) shows concentration of edges on immune-modulatory axes.

**Figure 4:**
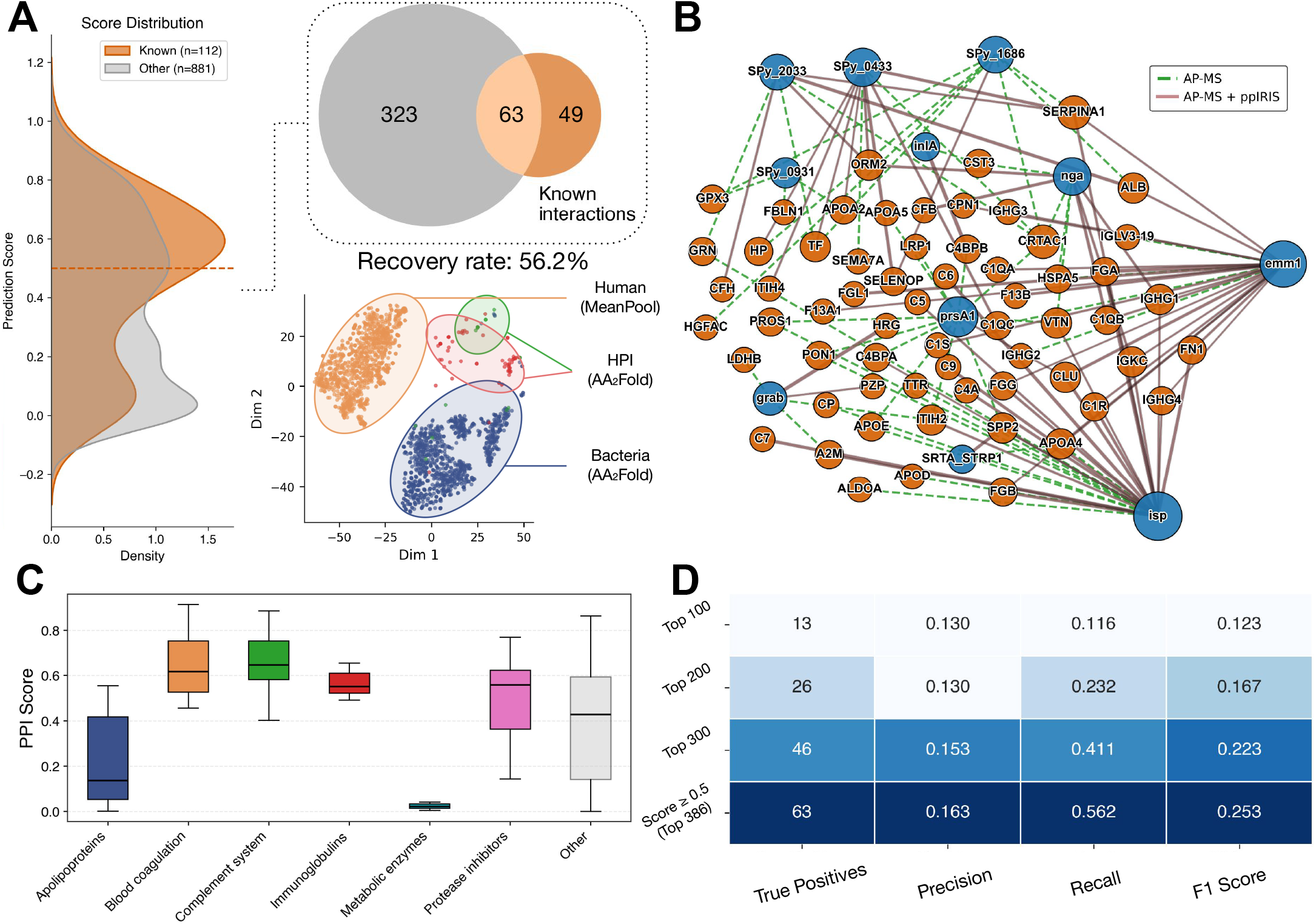
Cross-species PPI predictions between 16 GAS virulence factors and human plasma proteins. **A)** The distribution of ppIRIS prediction scores shows that known AP-MS interactions (orange) are systematically enriched at higher scores compared to all other candidate pairs (gray). A Venn diagram illustrates the overlap between the 386 predicted pairs at a score threshold of 0.5 and the 112 experimentally detected interactions, with 63 interactions recovered (56.2%). A tSNE plot of the embeddings reveals distinct clusters for bacterial (AA2Fold) and human (MeanPool) proteins, with host-pathogen interaction proteins (AA2Fold) enriched in a third cluster. **B)** A network representation highlights validated GAS-human interactions, with AP-MS interactions shown in dashed green and ppIRIS recovered interactions shown in red. **C)** Boxplots of prediction scores stratified by human protein class reveal that complement system, blood coagulation, and protease inhibitor proteins are consistently scored higher than apolipoproteins and metabolic enzymes. **D)** Cumulative performance across different prediction cutoffs, reporting the number of true positives, precision, recall, F1 score, and recovery rate. At the 0.5 threshold, ppIRIS recapitulates 63 of 112 known interactions with precision 0.163, recall 0.562, and F1 score 0.253.

Stratification of the predictions by human protein categories revealed systematic performance differences (**Figure 4C**). The highest prediction scores and recovery rates were observed for proteins of the complement system, blood coagulation cascade, and protease inhibitors. This is biologically plausible, since these protein families are known to engage directly with bacterial virulence factors in processes related to immune evasion and host defense [17]. In contrast, interactions involving apolipoproteins were poorly recovered, and for metabolic enzymes, the recovery was negligible. These categories were underrepresented in the training dataset and display sequence diversity that likely challenges feature transfer from intra-species PPIs. These results underscore that ppIRIS is biased toward well-represented classes of proteins in the training data, and that performance can be enhanced by expanding the training corpus to include additional examples of underrepresented categories. The precision remains modest, which is expected given the strong class imbalance (11.3% positives in the labeled subset), as well as the presence of likely false negatives among the pseudo-negatives. For triage use, cutoffs below 0.5 increase recall (Figure 4D) and could be adapted to experimental throughput. Taken together, these results demonstrate the generalizability of ppIRIS from intra-species to inter-species PPI prediction recovering a substantial fraction of experimentally validated GAS-human PPIs. The enrichment of validated interactions, the concentration of high-scoring predictions in immunologically relevant pathways, and the emergence of plausible novel candidates together support the utility of ppIRIS for cross-species prediction.

### Experimental validation of ppIRIS predictions

To validate whether ppIRIS shortlists are experimentally enriched, we performed two AP-MS assays (named as interactome A and B) using a bait-specific pulldown strategy from prey mixture incubation (Figure 5), scored with Mass Spectrometry Interaction STatistics (MiST) [30]. Interactome A was focused on host binding partners of two recombinant GAS M1 virulence factors-C5a peptidase (C5a) and streptolysin O (SLO) and purified via Strep-Tactin affinity. We have previously reported that the pathogenicity of SLO involves the conversion of plasminogen to plasmin [31], and now expanded on that knowledge. The prey space comprised nearly 400 identified and quantified human plasma and saliva proteins. Among ppIRIS-nominated pairs for both plasma and saliva proteins, 53% were supported by high-confidence AP-MS MiST score (above 0.75).

**Figure 5:**
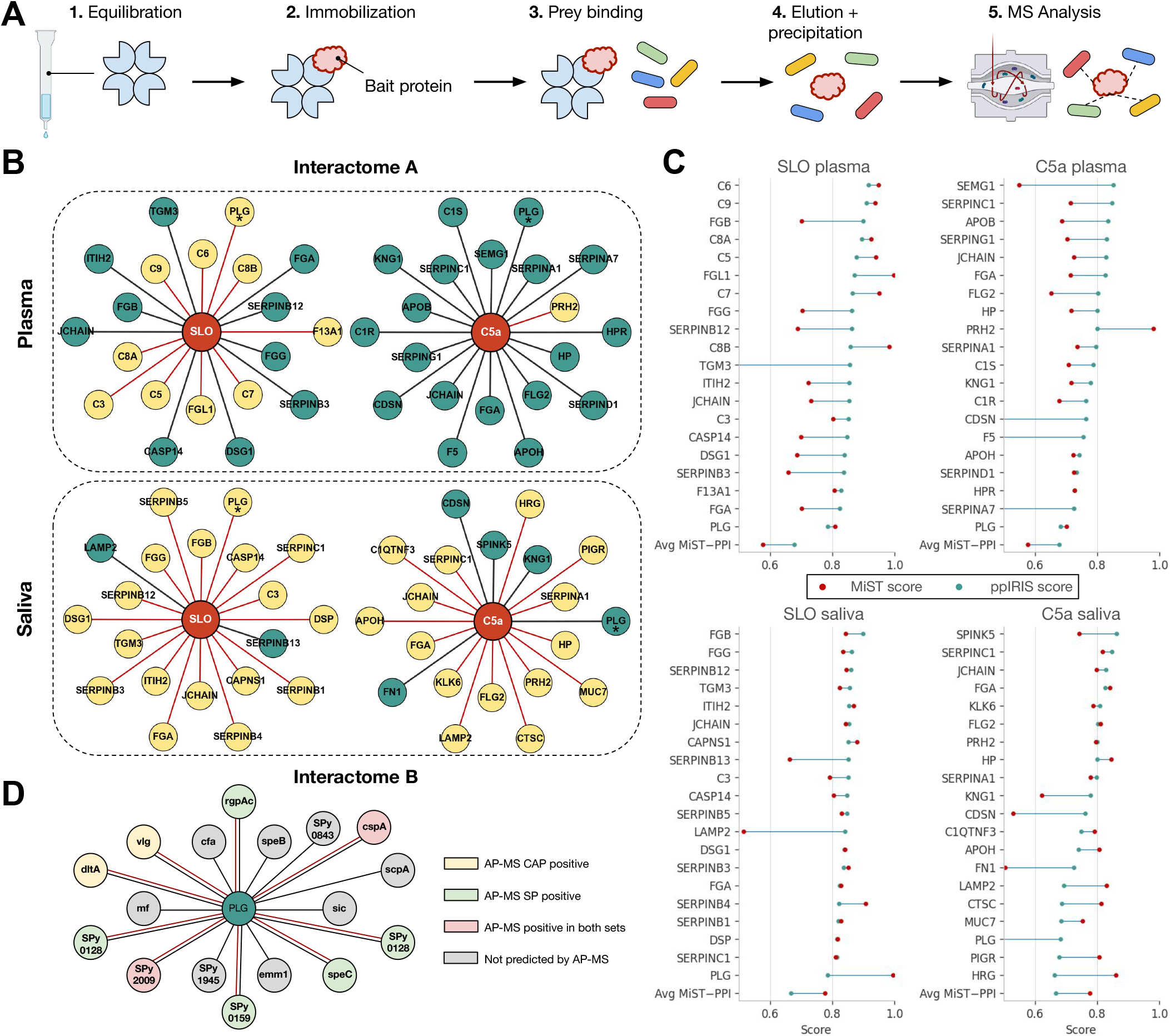
Prospective validation of ppIRIS predictions by AP-MS. **A)** Schematic of the AP-MS workflow. **B)** Top-20 predictions are shown for interactome A, where recombinant C5a peptidase and streptolysin O were used as baits against human plasma and saliva. Nodes highlight validated human proteins, with red edges denoting ppIRIS-predicted interactions confirmed by AP-MS. Only top-20 are illustrated except for PLG given its significance interaction with SLO [31]. **C)** The difference between the ppIRIS (green points) and the MiST (red points) scores for top 20 predictions from interactome A sorted by PPI score and grouped by bait and source. **D)** For interactome B, human plasminogen served as bait against GAS cell-wall-associated (CAP) and secreted proteins (SP) fractions. Prey nodes are colored by AP-MS evidence (CAP-positive, SP-positive, both, or not detected).

In a different setup with ECH-Lysine resin, interactome B assessed the interaction landscape of human plasminogen by GAS proteins. Roughly 100 GAS proteins were confidently quantified across the two fractions, and half (50%) of ppIRIS plasminogen-GAS predictions were experimentally confirmed. Supported interactions included known or plausible plasminogen-binding surface adhesins and secreted enzymes, alongside uncharacterized proteins. Across both assays ppIRIS thus produced conservative candidate sets with high prospective precision (0.53 and 0.50) despite large underlying search spaces. These results validate ppIRIS as an efficient framework to complement experimental effort for host-pathogen interactions.

### Towards an interaction landscape of GAS virulence factors with human proteins

We further explored the interaction landscape of GAS M1 serotype by predicting interactions across over thirty GAS virulence factors and thousands of human proteins (**Figure 6**). We leveraged the same set of human plasma and saliva proteins used in the previous section, as well as high-confidence human surfaceome proteins [32]. This analysis yielded thousands of predicted PPIs. We further filtered the list up to five interactions per GAS virulence factor for better visualization and interpretation. The resulting network revealed coherent functional modules that mirror the host environments encountered by GAS. Predicted edges concentrated on *(i)* complement and membrane-attack complex components consistent with innate immune lysis defense [33]; *(ii)* epithelial protease control centered on serpins and protease regulators, in line with GAS protease activity at mucosal surfaces; and *(iii)* barrier-fluid proteins abundant in saliva and plasma. These modules contained multiple convergent edges from distinct virulence factors, suggesting multi-protein targeting rather than isolated one-off contacts (**Figure 6B**).

**Figure 6:**
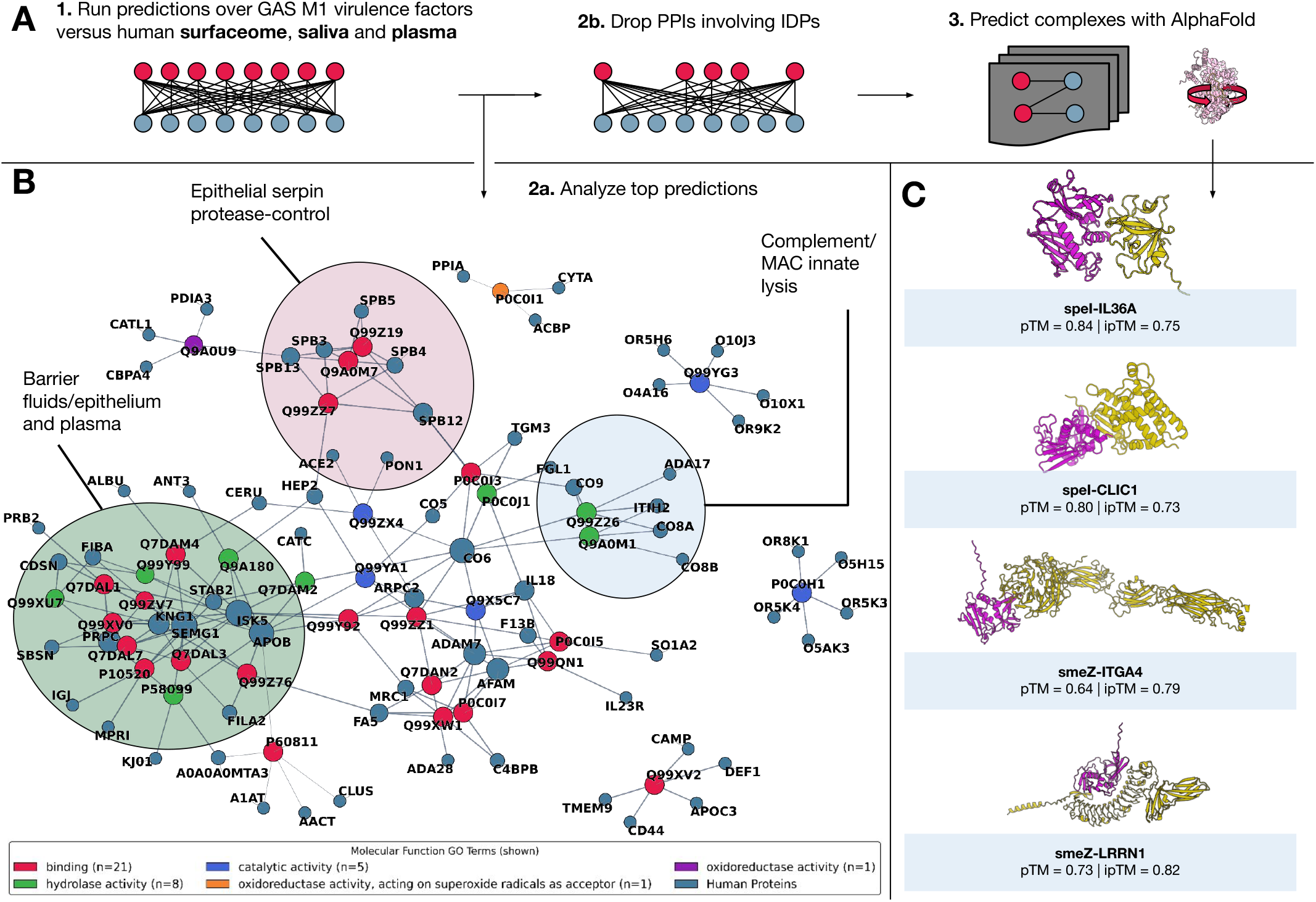
Predicted interaction landscape of GAS M1 virulence factors with human proteins. **A)** Schematic of the ppIRIS workflow: predictions were run against human surfaceome, saliva, and plasma proteins; pairs involving intrinsically disordered proteins were removed; and top candidates were forwarded to AlphaFoldmultimer for complex modeling. **B)** Network representation of top-scoring predictions, showing coherent functional modules targeted by multiple virulence factors, including complement/MAC components (innate lysis), epithelial serpin-mediated protease control, and barrier-fluid proteins. Node colors correspond to molecular function categories (binding, catalytic, hydrolase, oxidoreductase). For better visualization, we limited interactions to the top 5 interactions per GAS protein. **C)** Example AlphaFold-multimer models of selected GAS-human complexes. Four representative predictions are shown: SpeI-IL36A (pTM = 0.84, ipTM = 0.75), SpeI-CLIC1 (pTM = 0.80, ipTM = 0.73), SmeZ-ITGA4 (pTM = 0.64, ipTM = 0.79), and SmeZ-LRRN1 (pTM = 0.73, ipTM = 0.82). These span a range of biological plausibility, from well-supported interactions (e.g., SmeZ-ITGA4) to more speculative candidates requiring experimental validation.

We further structurally assessed selected high-ranked pairs with AlphaFold-multimer. We filtered our predictions to drop PPIs involving any protein that was at least 50% disordered. While this decreased the number of predicted PPIs, given the extensive number of intrinsically disordered proteins in the GAS proteome, this allowed for structure-based validation. Four representative complexes are shown in **Figure 6C** including SpeI-IL36A, SpeI-CLIC1, SmeZ-ITGA4, SmeZ-LRRN1. The SpeI-IL36A interaction is more speculative but biologically conceivable given that IL-36 cytokines can be released and proteolytically activated during pathogen-induced tissue damage [34]. A putative SpeI-CLIC1 interaction is also detected, given the involvement of CLIC1 in NLRP3 inflammasome activation that can take place during microbial infections [35]. The SmeZ-ITGA4 aligns with known interactions of pathogens with integrins for adhesion and uptake. Also involving the same GAS superantigen, SmeZ-LRRN1 is also detected, as an interaction between neuronal assembly protein LRRN1 and bacterial antigens have not been previously described. However, leucine rich repeat-containing proteins are known to be involved in the detection of pathogen molecular patterns [36], and GAS is known to infiltrate and interact with CNS cells, including sensory neurons [37, 38], making this an intriguing candidate for future validation.

## Discussion

ppIRIS provides a fast, sequence-based framework for large-scale bacterial and host-pathogen PPI prediction by fusing complementary evolutionary (ESM-C 300M) and structure-aware (ProstT5 3Di) embeddings within a lightweight Siamese architecture. Rather than introducing deeper token-level transformers at inference, we show that a compact interaction head together with pooling can reach or exceed the F1 of substantially heavier baselines (e.g., D-SCRIPT, TUnA) [5, 6, 9] while keeping training and inference times minimal. The main technical contributions are: *(i)* simple dual-embedding fusion that improves recall without sacrificing precision, *(ii)* a “domain shift” strategy for pooling (AA2Fold vs. mean pooling for ProstT5 3Di) during training followed by unification at inference to encourage invariance to pooling-induced perturbations, and a compact feature interaction module that augments raw pair features without expensive residue-level cross-encoders. Together these design decisions emphasize practical deployability (rapid proteome scans on more broadly available GPUs) and enable iterative experimental cycles, complementing recent calls for lightweight PPI predictors [9]. Complementarity of embeddings was empirically supported. ESM-C alone yielded the highest precision, whereas adding ProstT5 3Di boosted recall and overall F1, indicating that coarse structural context rescues true positives missed by purely evolutionary signals. Practically, a moderate 5:1 ratio maximized structural signal while avoiding the recall collapse and potential precision inflation artifacts associated with more aggressive negative pairing, consistent with commonly used negative:positive ratios in sequence-based PPI modeling [39].

The biological application illustrated three primary use cases. (1) Intra-proteomic scanning of the GAS M1 serotype rapidly highlighted under-annotated functional clusters (transport, stress, metal scavenging, phage-related modules) beyond well-known ribosomal assemblies. (2) Cross-species screening recovered 56.2% of curated GAS-human plasma PPIs at a 0.5 threshold despite the model never receiving cross-species supervision, with enrichment in complement, coagulation, and protease inhibition pathways consistent with immune evasion biology [17]. (3) Expansion to virulence factor-human surfaceome interactions revealed convergent multi-targeting of adhesion (integrin-related) and barrier-associated proteins. AlphaFold-multimer assessment resulted from plausible interfaces (e.g., SmeZ-ITGA4) to speculative cytokine or neuronal candidates (e.g., SmeZ-LRRN1, and SpeI-IL36A). The AP-MS validation also indicated that ppIRIS ranked predictions were enriched (precision 0.53 and 0.50 across two interactomes) with expected low recall under sparse sampling, in line with prior plasma/saliva interaction mapping efforts [3, 17, 40]. This supports the usage of ppIRIS when studying cross-species PPIs by deploying the model to reduce the experimental search space rather than to exhaustively consider all interactions.

Failure modes and bias were also evident. Recovery was weaker for apolipoproteins and metabolic enzymes, families underrepresented in training and more sequence-diverse, implying a need for targeted augmentation or domain adaptation. Residual species/family performance differences may partly reflect differential evolutionary coverage in pretrained language models rather than intrinsic interaction propensities. Negative sampling (random non-documented pairs) risks label noise and can inflate precision if class priors are misaligned with biological prevalence [9, 39]; prior work has highlighted evaluation leakage and sampling pitfalls [41]. Future work should explore different strategies for generating negative examples, functional stratified sampling, or semi-supervised pseudo-label refinement.

Structural post-filtering must be interpreted cautiously. Low AlphaFold-multimer confidence for bacterial or intrinsically disordered partners does not conclusively disprove interaction, while high pTM/ipTM may still reflect docking artifacts [25, 29, 42]. Incorporating disorder-aware or coarse-grained interface plausibility metrics and uncertainty estimates such as Monte Carlo dropout, deep ensembles, or spectral-normalized Gaussian processes could improve decision calibration.

## Data and Code Availability

The ppIRIS codebase, including the dataset preprocessing, embedding extraction, training, and inference pipelines, is available at https://github.com/lupiochi/ppiris. All AP-MS and related proteomics datasets deposited to PRIDE open-access repository will be made publicly available.

## Acknowledgment

HK was supported by the French Agence Nationale de la Recherche (ANR), under grants ANR-22-CPJ2-0075-01, and ANR-24-CE45-4243-01.

## Contributions

LP, YK, and HK conceived the idea. LP developed the deep learning model in discussion with YK and HK, and performed the data analysis. DT, and JM performed and provided experimental data. HK and YK supervised the project. LP and HK wrote the initial draft with input from all authors.

## Competing interests

The authors declare that they have no conflict of interest.

## Materials & Methods

### Data Collection

#### Bacterial PPIs

Protein-protein interaction (PPI) data from gram-positive bacteria were retrieved from STRING DB v12. True-positive (TP) interactions were defined by filtering entries with an experimental score *>* 600 and combined score *>* 990 as positives [2]. True-negative (TN) interactions were generated by random sampling of protein pairs not reported to interact in STRING. We used negative:positive ratios of 5:1 and 10:1 in separate experiments, following established practices in PPI prediction [39]. The GAS M1 serotype proteome was obtained from UniProt (UP000000750) and matched to STRING protein identifiers by BLAST, requiring over 95% identity and 90% coverage, yielding ∼800 matched PPIs. If the function was unavailable in UniProt, DeepGO2 [24] was used to predict the GO terms for molecular function.

#### Human PPIs and cross-species ground truth

Human PPIs were taken from the Bernett et al. [20] dataset. PPI datasets for other species, including *D. melanogaster, C. elegans, S. cerevisiae* and *E. coli*, were obtained from Sledzieski et al. [5]. For host-pathogen evaluation, we used the curated panel from our previous work [17], comprising 16 well-characterized GAS M1 virulence factors and 62 abundant human plasma proteins with 112 AP-MS-validated interactions.

#### Bacterial taxonomic data

An advanced search and curation was performed on the BacDive database [43] to identify gram-positive bacteria with available PPI data in STRING. The search was filtered to include only entries with the following criteria: (i) Gram stain = “positive”, and (ii) Type strain = “true”. The search results were merged with metadata from STRING, which yielded a total of 794 unique taxon ids.

### Data Preprocessing

To mitigate information leakage, we created train/validation splits under sequence-similarity constraints using MMseqs2-based clustering [44]. To look for cross-split homology, we applied a pairwise search filter, removing any validation protein with *>* 40% similarity to a training protein (as suggested by other studies [5, 20]). Self-loops were excluded and duplicate pairs removed, and PPIs were only retained if both partners fell within the same split. For each target negative:positive ratio (5:1 and 10:1), we sampled negatives within source (human-human and bacterial-bacterial) while ensuring no overlap with positives. All splits were written with columns `protein1, protein2, label` and used consistently for training and evaluation. For cross-validation experiments, we additionally prepared a combined split with all proteins from gram-positive bacteria and humans using the same MMSeqs2-based clustering approach.

### Embedding extraction, pooling strategies, and cross-species bridging

#### Encoders and per-protein embeddings

We represented each protein sequence with the concatenation of two pretrained encoders: ESM-C 300M and ProstT5 3Di. For ESM-C, we first obtain token-level embeddings 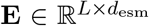 and compute a sequence embedding via attention pooling. 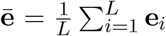 be the context vector. We score tokens by 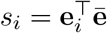, derive weights 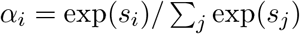, and set:

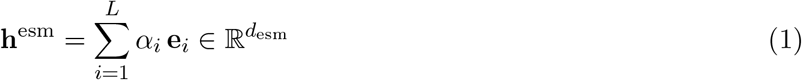

For ProstT5, we tokenized sequences as “`<AA2Fold> A A A`…” and compute either (i) the hidden state at the special prefix token (“AA2Fold”) as a CLS-like embedding, 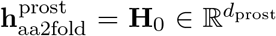 the mean of amino-acid token states (excluding the AA2Fold prefix), 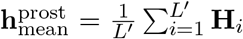. The outputs of the two encoders are fused by concatenation on the set of protein IDs common to both extractors, and fused vectors are stored in HDF5 with associated IDs and metadata, as:

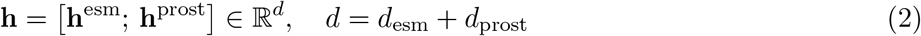

#### Pooling-based domain shift at training, unification at inference

Let *x* ∈ 𝒳 be a protein. We formed per-protein embeddings by concatenating the base attention-pooled ESM-C embeddings with a ProstT5-3Di:

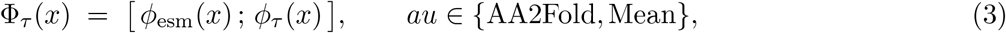

where *ϕ*_AA2Fold_ is the `<AA2Fold>` token pooling and *ϕ*_Mean_ is residue-wise mean pooling.

During training we intentionally used different poolings per species: bacteria → `AA2Fold`, human → `Mean`. If 𝒟_B_ and 𝒟_H_ denote the labeled pair distributions for bacteria and human, we minimized:

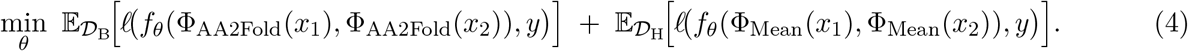

Because `AA2Fold` and `Mean` emphasize different aspects of the ProstT5-3Di representation, this created a domain shift in the inputs seen by the head while preserving labels. The shared-weights head was therefore pushed to focus on interaction cues that survived a change of pooling and to ignore pooling-specific artifacts.

During inference, however, we unified the readout. We chose one pooling *ψ* ∈ {AA2Fold, Mean} and applied the same Φ_*ψ*_ to both protein sets before scoring. Here, we proceeded with the AA2Fold pooling as our default choice. This removes the explicit species cue carried by the pooling itself and puts host and pathogen in the same representation space, which matches the invariance encouraged during training.

We relied only on a simple smoothness intuition: the scoring head changes its output gradually, as small changes in an input embedding produce small changes in the predicted interaction score. We defined the per-protein pooling gap as:

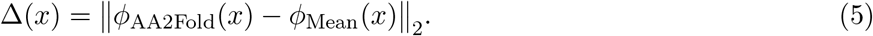

Swapping the pooling of just one protein in a pair adjusts its embedding by Δ(*x*), so the score shifts by a constant multiple of that change. Swapping both poolings therefore contributes additively:

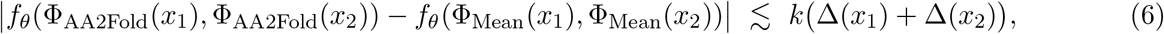

For some modest sensitivity constant *k* observed in practice, i.e., not explicitly enforced. Training with both poolings encourages the model to depend on interaction signals stable under this swap (keeping Δ’s effect small); using a single unified pooling at inference sets Δ(*x*) = 0 and removes this variation source entirely.

### Algorithm

#### Model

ppIRIS is a Siamese dual-branch network operating on fused per-sequence embeddings (input dim *d* = 1984). Each branch applies a shared MLP encoder with two fully connected blocks (defaults: hidden 512, output 256), each followed by batch normalization and ReLU; dropout (*p* = 0.5) is applied to intermediate blocks. To let each branch attend to its partner, we used a MultiheadAttention layer [45] (2 heads) over the pair of encoded representations, producing two cross-attended vectors. Although classical attention gains most of its expressivity from token-level alignment, here we deliberately applied it at the (already pooled) sequence level for two reasons: (i) it implements a light-weight, parameter-efficient learned mixture of bilinear similarity functions softmax 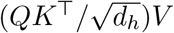 that adapts to heterogeneous embedding subspaces contributed by ESM-C vs ProstT5, something that static element-wise operations cannot re-weight dynamically; and multihead projections act as a low-rank factorization of an implicit full bilinear form between the two sequence embeddings, increasing relational capacity without incurring the *O*(*d*^2^) parameters of a naive bilinear layer. We then formed six features by concatenation: {rep_1_, rep_2_, cross_1_, cross_2_, |rep_1_ − rep_2_|, rep_1_ ⊙ rep_2_}, and passed them to a “combiner” MLP (512 hidden, BN+ReLU+dropout) ending in a spectrally-normalized linear unit that outputs a single logit interaction score.

#### Training

ppIRIS is optimized during training using the Adam optimizer with weight decay regularization and binary cross entropy loss. The best-performing model was determined by monitoring the validation loss. Gradient clipping techniques were also utilized to ensure stable training when necessary. The best set of hyperparameters was determined through a grid search.

### Benchmarks

We evaluated ppIRIS on the standardized PPI model benchmark from [9], following the same dataset partitions and protocol to ensure comparability with published baselines. Performance metrics (ROC-AUC, average precision, accuracy, specificity, precision, recall, balanced accuracy, F1, MCC) were computed on the validation/test splits. All results were obtained at a probability threshold of 0.5.

Let TP, FP, TN, FN denote true positives, false positives, true negatives, and false negatives at decision threshold *τ* = 0.5. Then:

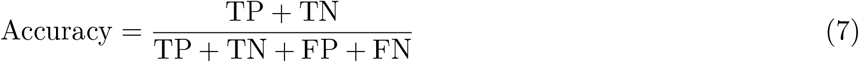

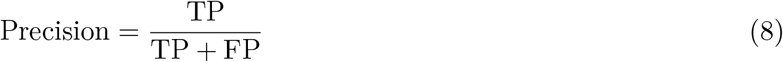

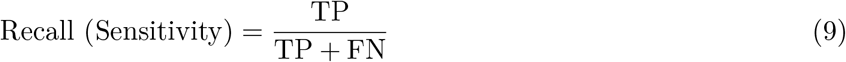

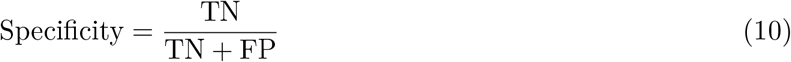

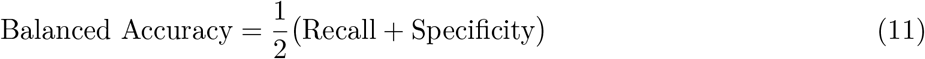

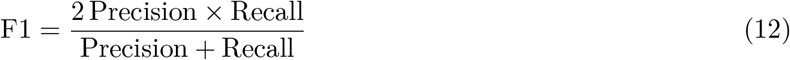

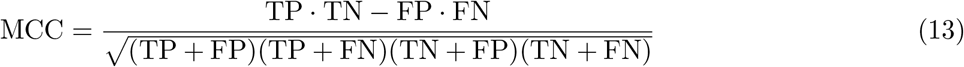

For threshold-free metrics, let scores induce an ordering of *N* pairs. Define at rank *k* the precision *P*_*k*_ and recall *R*_*k*_.

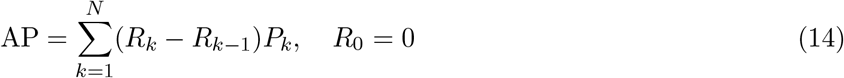

Receiver Operating Characteristic (ROC) curves plot True Positive Rate (TPR) versus False Positive Rate (FPR):

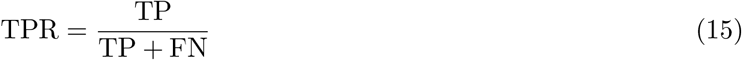

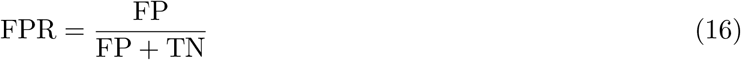

AUROC is the area under the ROC curve. AUPR is the area under the Precision-Recall curve.

Data from D-SCRIPT, TUnA and Topsy-Turvy when assessed using the D-SCRIPT multispecies datasets were obtained from their original manuscripts [5, 6, 22].

### Modelling and Evaluation of Top Predictions

To structurally assess high-confidence predictions, we selected top ppIRIS-scored pairs and generated models with AlphaFold-multimer v3 with default settings. Models were scored based on their pTM and iPTM values, with higher scores indicating better predicted structural quality.

### Affinity-purification assay

The affinity-purification mass spectrometry experiment was conducted as previously described [31]. Briefly, for interactome A, two bait GAS proteins together with a control GFP protein fused to affinity tags were designed, recombinantly expressed in *E. coli*, and purified. Pooled human plasma and pooled human saliva were sourced from Innovative Research. A 1 x PBS buffer prepared from tablets (Sigma-Aldrich) was used to dilute the plasma prey to 50 % (v/v). Pooled saliva was centrifuged and filtered before use as prey input. Bait proteins were first immobilised on pre-washed Strep-Tactin Sepharose resin (IBA Lifesciences GmbH) and then washed repeatedly with PBS to remove unspecific bindings. Protein complexes were eluted with freshly made biotin solution (Sigma-Aldrich); the pull-down samples were precipitated with trichloroacetic acid (TCA), repeatedly washed with cold acetone, concentrate to full dryness in a SpeedVac (Eppendorf), and the dried pellets were reconstituted in 100 mM ammonium bicarbonate for downstream in-solution digestion. Three independent experiments were performed per bait-prey condition in interactome A. For interactome B, native human plasminogen was purchased from Sigma-Aldrich. Prey mixtures were prepared separately from overnight cultures of M1 strain *Streptococcus pyogenes*. The bacterial medium (Sigma-Aldrich) was centrifuged, and the supernatants were collected, concentrated, and designated the secreted-protein fraction. Pellets were resuspended in phenylmethylsulphonyl fluoride buffer, supplemented with freshly prepared mutanolysin buffer, and incubated in a ThermoMixer (Eppendorf); after centrifugation, the supernatants were collected and designated the cell-wall-associated protein fraction. Plasminogen bait was incubated with pre-equilibrated ECH-lysine bead slurry (Cytiva) for immobilisation, after which the prey mixture from either fraction was loaded onto the column. The column was washed with 50 mM sodium dihydrogen phosphate containing 0.1 M sodium chloride to remove the unspecific bindings and, finally, the formed protein complexes were eluted with 0.1 M sodium chloride supplemented with 50 mM *ε*-aminocaproic acid. The same TCA precipitation and protein concentration was applied to the elution as described above.

### Mass spectrometry sample preparation and analysis

Routine in-solution digestion was performed on both GAS protein fractions and the various pull-down samples. Urea (Sigma-Aldrich) and tris(2-carboxyethyl)phosphine (Thermo Fisher Scientific) were used for protein denaturation and disulfide reduction, followed by alkylation with iodoacetamide (Sigma-Aldrich). Proteins were digested overnight with 1:20 w/w trypsin (Promega), and peptides were cleaned up using a C18 spin column (Thermo Fisher Scientific). For interactome B, portions of both the secreted-protein and cell-wall-associated protein fractions underwent extra fractionation with the High-pH Fractionation Kit (Thermo Fisher Scientific) to improve identification of low-abundance species. Eluted peptides were dried in a SpeedVac (Eppendorf) and reconstituted in buffer A (2 % acetonitrile, 0.2 % formic acid; Thermo Fisher Scientific) for mass spectrometric analysis. For interactome A, peptides were loaded onto an EASY-nLC 1200 system coupled to a Q Exactive HF-X hybrid quadrupole-Orbitrap mass spectrometer (Thermo Fisher Scientific). Each sample was injected once for data-independent acquisition (DIA). Liquid-chromatography separation comprised solvent A (0.1 % formic acid) and solvent B (0.1 formic acid in 80 % acetonitrile). A linear gradient from 3 % to 38 % solvent B was run over 90 min at 350 nL/min. MS data acquisition began with an MS1 scan (390 to1210 m/z) at 60,000 resolution, AGC target 3e6, and maximum injection time 100 ms, followed by MS2 scans with a fixed 26.0 m/z isolation window, 30,000 resolution, AGC target 1e6, injection time 120 ms, and normalised collision energy (NCE) of 30. The same LC-MS set-up was used for interactome B. For the GAS protein fractionation samples, an additional data-dependent acquisition (DDA) method was applied: a gradient from 5 % to 38 % solvent B over 120 min, MS1 scan (350-1650 m/z) at 60,000 resolution, AGC 3e6, and 50 ms injection time, followed by the top 20 MS2 scans at 15,000 resolution, AGC 1e5, 25 ms injection time, and NCE of 27. For DIA acquisition in interactome B, the same gradient described above was used for 90 min period. MS1 scans covered 350-1650 m/z at 120,000 resolution, AGC 3e6, and 60 ms injection time, followed by 44 MS2 scans at 30,000 resolution, AGC 3e6, and stepped NCE comprised of 25.5, 27, and 30.

### Quantitative proteomics and interaction analysis

For library construction, a predicted spectral library was generated from proteome UP000005640 for interactome A. For interactome B, AE-MS DIA data and fractionation DDA data were combined to build a hybrid spectral library using FragPipe [46] with proteome UP000000750. DIA-NN was employed to search the MS data and generate the quantification matrix; inter-run normalisation was disabled, consistent with the bait-specific affinity-purification design. A bait-prey matrix was then constructed for each condition and scored using the MiST (Mass Spectrometry Interaction STatistics) workflow. Confident bait-prey interactions were experimentally defined as those with a MiST score *>*0.75, derived from the combined specificity, abundance, and reproducibility metrics.

## Notes

### Competing Interest Statement

The authors have declared no competing interest.

