## Supplementary Material for "ppIRIS: deep learning for proteome-wide prediction of bacterial protein-protein interactions"

#### Supplementary Figures

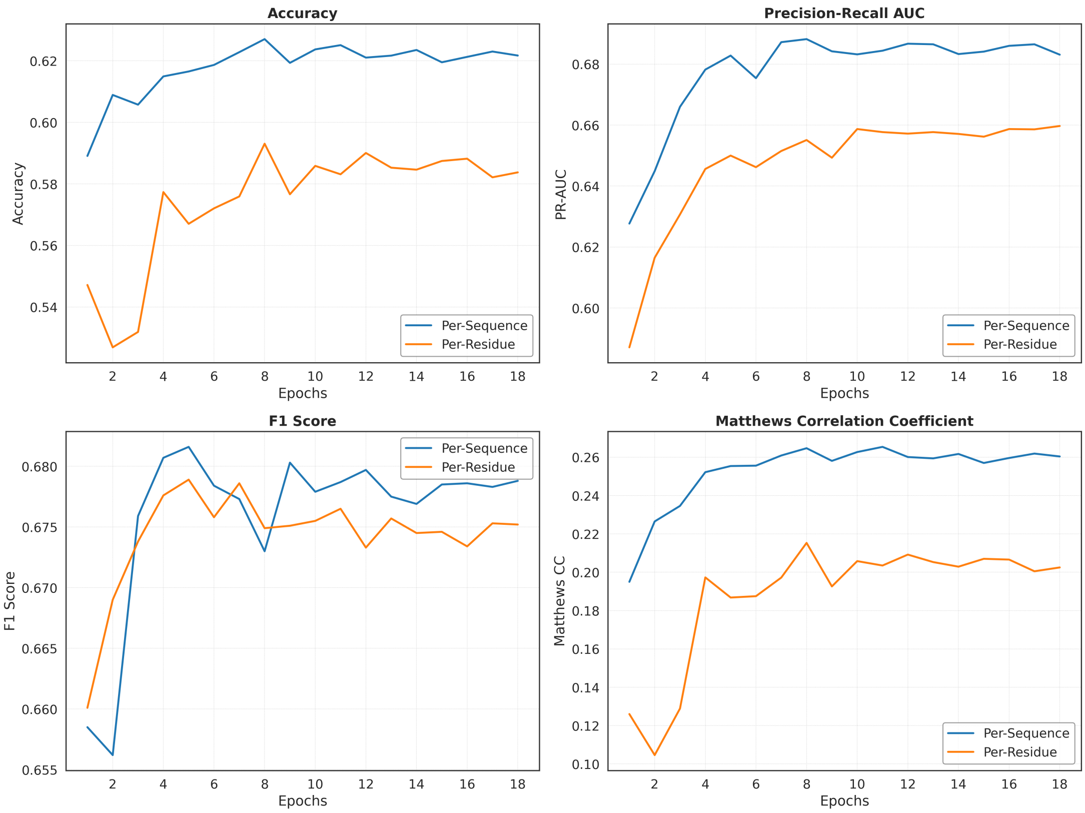


Figure S1: Comparison of performance metrics when using per-sequence versus per-token embeddings on the human gold-standard PPI dataset. While per-token embeddings theoretically capture positional information, our results show they provide minimal additional benefit for PPI prediction while substantially increasing computational demands. We therefore used per-sequence embeddings throughout our study for efficiency.

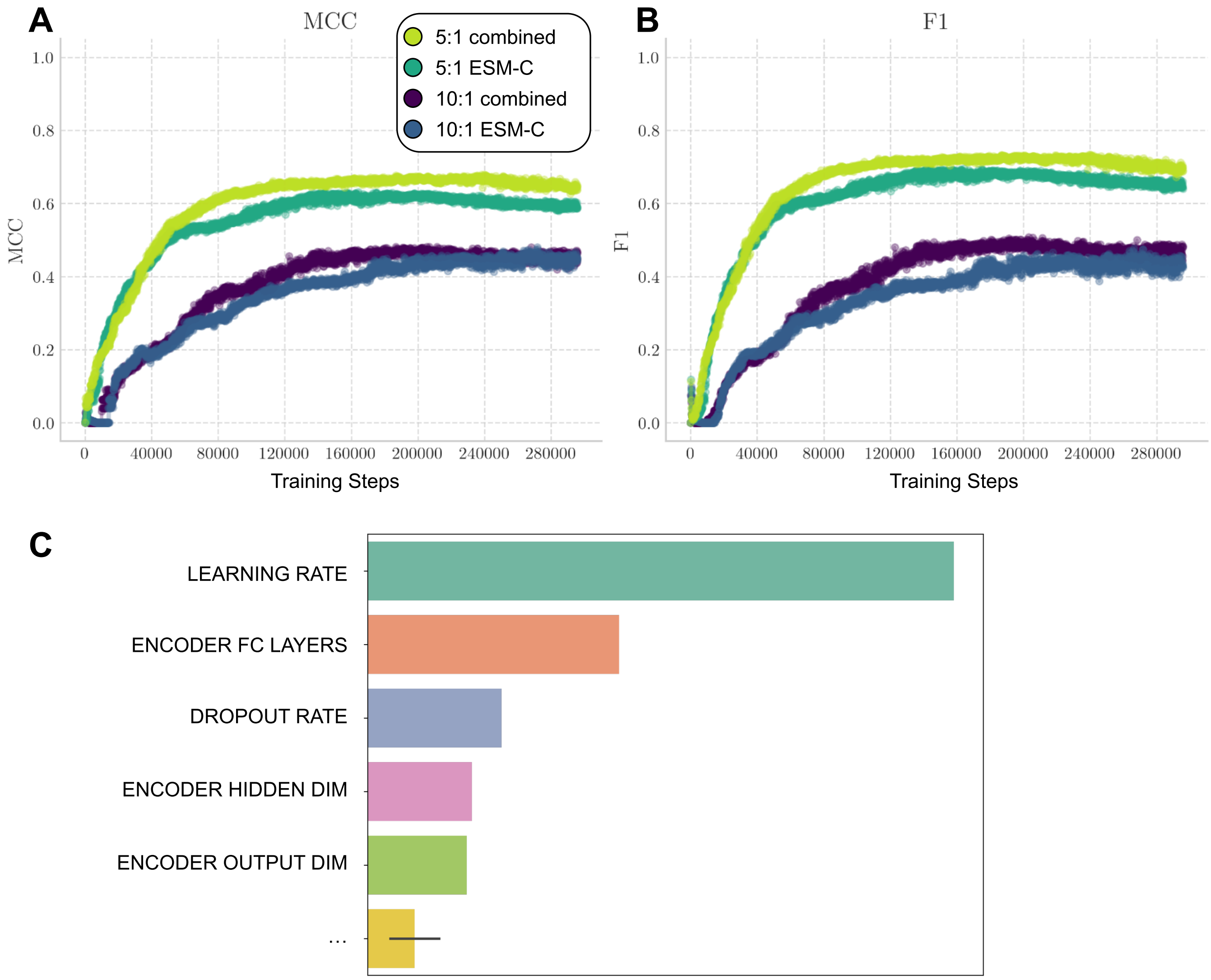


Figure S2: Prediction results for the top 10,000 intra-proteomic interactions based on the predicted PPI score. **A-B)** Validation MCC and F1 score per training step. Values in blue and purple represent the metrics for the 0:1 split using only ESM-C 300M embeddings and 10:1 split using the combined embeddings, respectively. Values in dark green are for the 5:1 ESM-only dataset, whereas light green are for the 5:1 using the combined embeddings. It becomes evident that the model is able to achieve a high F1 score and MCC, with the combined embeddings achieving the best performance. **C)** Hyperparameter importance on the classification performance, wherein the most important hyperparameters are the learning rate and the number of hidden layers in the encoder, followed by the dropout rate.

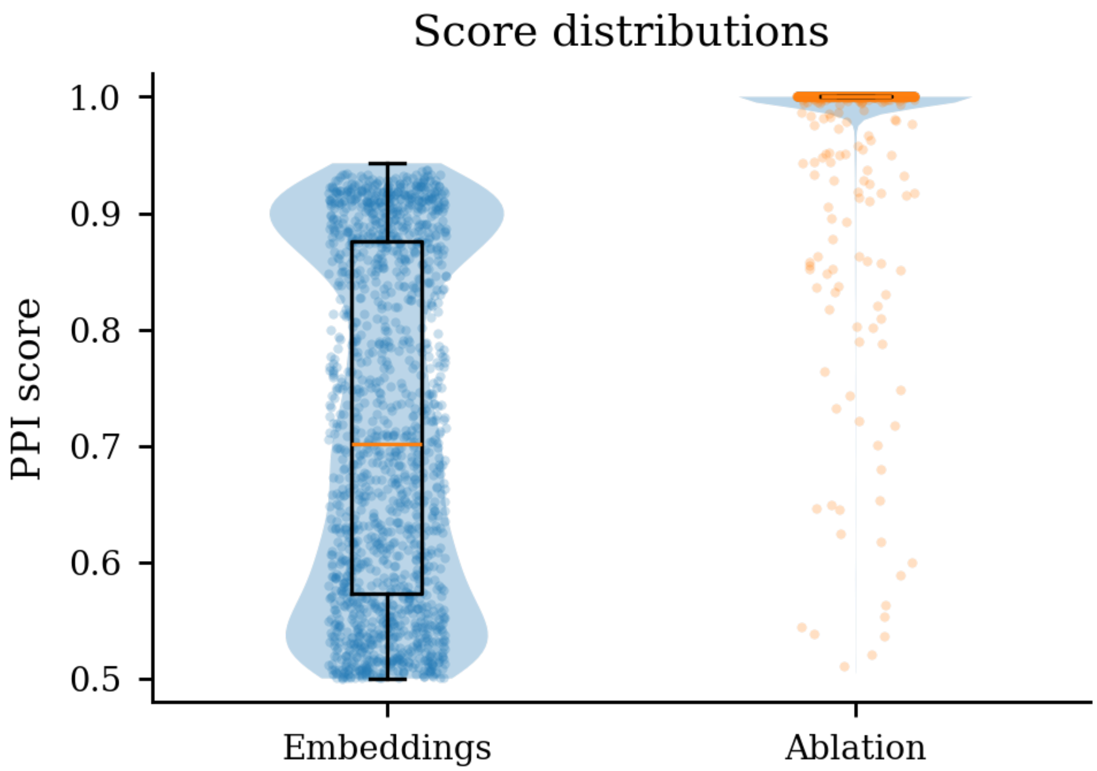


Figure S3: Score distributions using true embeddings versus random noise, for predictions with a PPI interaction score over 0.5. The model with true embeddings (blue) shows a clear distribution of predictions across several values, while the model with random noise (orange) simply predicted everything with a high score, indicating no meaningful discrimination.

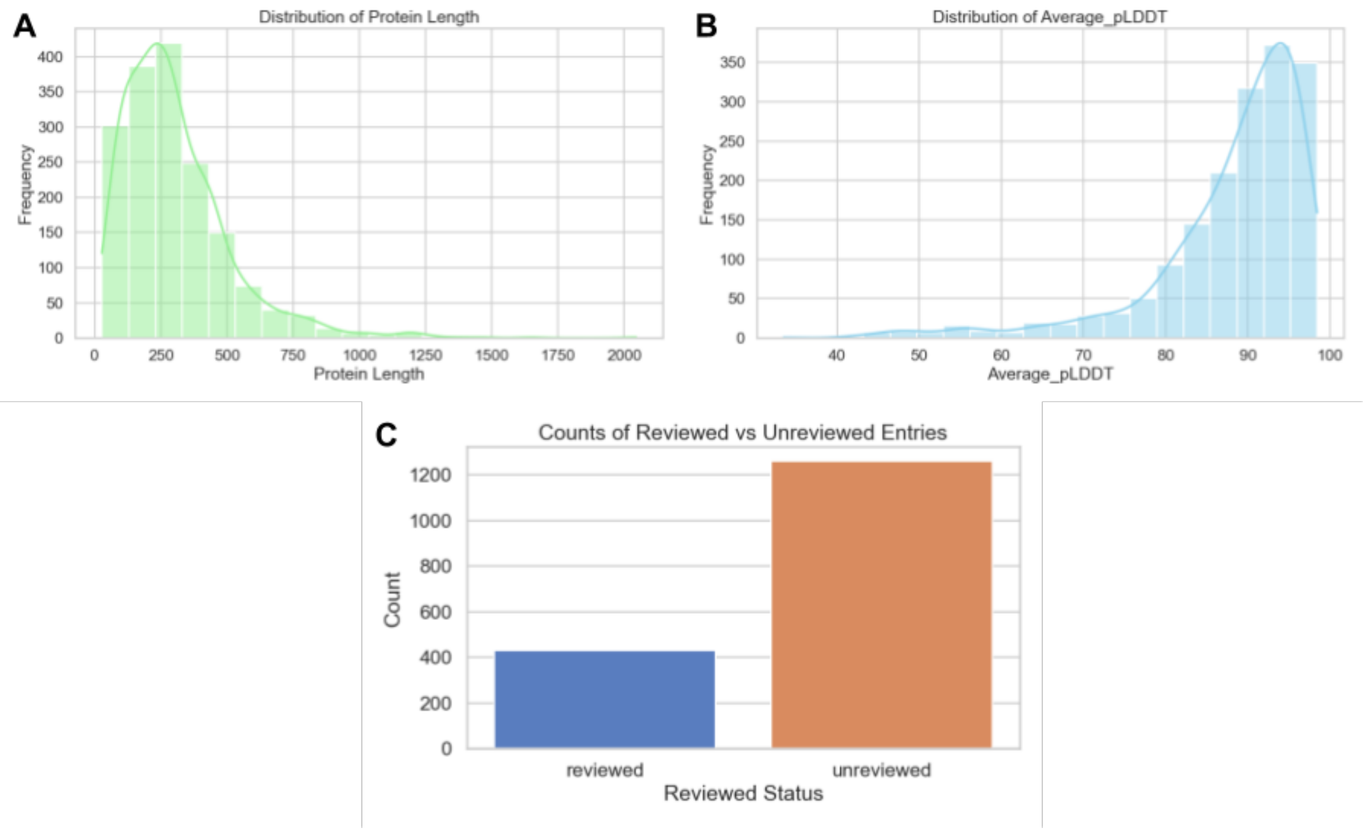


Figure S4: Characteristics of the GAS M1 Serotype proteome used for ppIRIS predictions. A) Protein length distribution showing most proteins between 100-500 amino acids. B) Distribution of average pLDDT scores across the proteome, reflecting confidence in structural predictions. C) Proportion of reviewed (manually curated) and unreviewed proteins in the proteome.

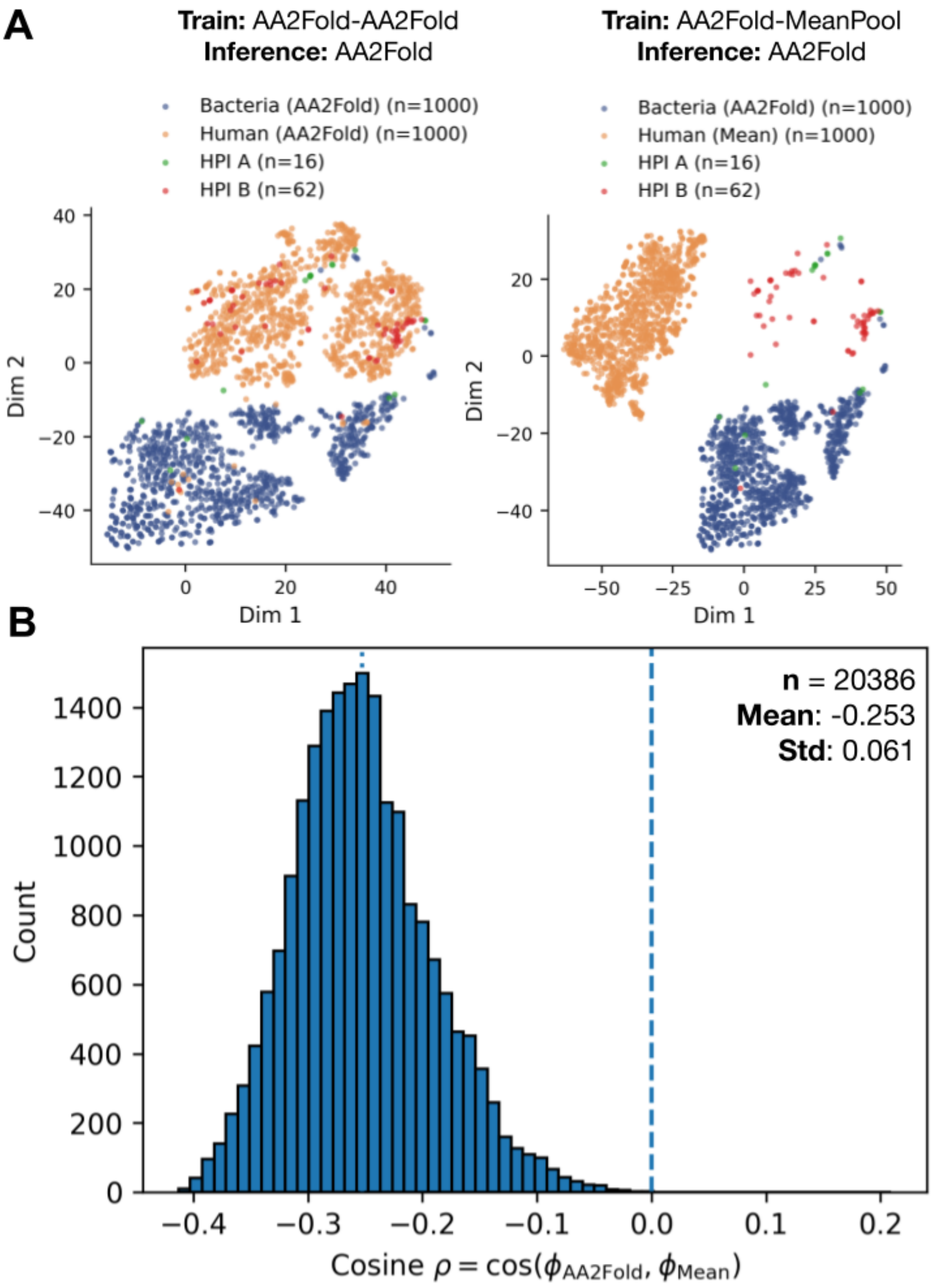


Figure S5: Intentional domain shift in cross-species predictions using different pooling strategies for ProstT5 embeddings. A) Two-dimensional tSNE projection of protein embeddings. On the left, both bacterial and human embeddings with AA2Fold-based pooling. On the right, bacterial embeddings with AA2Fold-based pooling and human embeddings with a mean pooling strategy. In both cases, the host-pathogen protein sequence embeddings were pooled with the AA2fold approach, showing a better separation from the rest when human and bacterial proteins from the training set are represented with different embeddings. B) Cosine similarity histogram between AA2Fold and MeanPool embeddings for matched proteins is broadly negative, supporting a notable difference between per-sequence embeddings obtained through different pooling approaches.

### Supplementary Tables

Table S1: Cross-species PPI performance (AUROC, AUPR) comparing ppIRIS with prior sequence-based models (including Topsy-Turvy). ppIRIS uses a 0.5 decision threshold. Human results are 5-fold CV means on the D-SCRIPT dataset. The performance metrics for Topsy-Turvy, PIPR, D-SCRIPT and TUnA for all species are reported in the Topsy Turvy and TUnA manuscripts.

| **Species** | **Model** | **AUROC** | **AUPR** |
|---|---|---|---|
| *D. melanogaster* | PIPR | 0.728 | 0.278 |
| | D-SCRIPT | 0.890 | 0.605 |
| | TUnA | 0.971 | 0.855 |
| | Topsy-Turvy | 0.921 | 0.713 |
| | **ppIRIS** | **0.976** | **0.876** |
| *C. elegans* | PIPR | 0.757 | 0.346 |
| | D-SCRIPT | 0.853 | 0.550 |
| | TUnA | 0.960 | 0.834 |
| | Topsy-Turvy | 0.906 | 0.700 |
| | **ppIRIS** | **0.973** | **0.866** |
| *S. cerevisiae* | PIPR | 0.718 | 0.230 |
| | D-SCRIPT | 0.790 | 0.399 |
| | TUnA | 0.898 | 0.641 |
| | Topsy-Turvy | 0.850 | 0.534 |
| | **ppIRIS** | **0.925** | **0.696** |
| *E. coli* | PIPR | 0.675 | 0.271 |
| | D-SCRIPT | 0.770 | 0.513 |
| | TUnA | 0.883 | 0.677 |
| | Topsy-Turvy | 0.805 | 0.556 |
| | **ppIRIS** | **0.912** | **0.747** |

Table S2: Comparison of General Model Performance Metrics for Different Negative-to-Positive Ratios and Embedding Groups. All datasets were clustered at 40% sequence identity threshold.

| **Embeddings** | **Split** | **W. ROC AUC** | **Avg. Prec.** | **Prec.** | **Recall** | **F1** | **MCC** |
|---|---|---|---|---|---|---|---|
| ESM-C | **5:1** | 0.9401 | 0.9226 | **0.7135** | 0.6193 | 0.6631 | 0.6035 |
| | **10:1** | 0.9389 | 0.9216 | 0.6575 | 0.3329 | 0.6242 | 0.4420 |
| ProstT5 3Di | **5:1** | 0.8436 | 0.8141 | 0.6083 | 0.2847 | 0.3879 | 0.3447 |
| | **10:1** | 0.8204 | 0.7834 | 0.4217 | 0.0982 | 0.1593 | 0.1693 |
| Combined | **5:1** | **0.9419** | **0.9248** | 0.6998 | **0.6807** | **0.6901** | **0.6292** |
| | **10:1** | 0.9392 | 0.9235 | 0.6641 | 0.4203 | 0.5148 | 0.4926 |

Table S3: Molecular function Gene Ontology (GO) term distribution among top 3,000 high-confidence ppIRIS predictions within the GAS M1 proteome after ribosomal proteins were removed. The table lists terms (abbreviated IDs), their definitions, and counts of associated proteins.

| **GO term** | **Definition** | **Count** |
|---|---|---|
| GO:0005488 | The selective, non-covalent, often stoichiometric, interaction of a molecule with one or more specific sites on another molecule. | 418 |
| GO:0003824 | Catalysis of a biochemical reaction at physiological temperatures. In biologically catalyzed reactions, the reactants are known as substrates, and the catalysts are naturally occurring macromolecular substances known as enzymes. Enzymes possess specific binding sites for substrates, and are usually composed wholly or largely of protein, but RNA that has catalytic activity (ribozyme) is often also regarded as enzymatic. | 225 |
| GO:0022857 | Enables the transfer of a substance, usually a specific substance or a group of related substances, from one side of a membrane to the other. | 149 |
| GO:0016787 | Catalysis of the hydrolysis of various bonds, e.g. C-O, C-N, C-C, phosphoric anhydride bonds, etc. | 45 |
| GO:0140640 | Catalytic activity that acts to modify a nucleic acid. | 18 |
| GO:0016740 | Catalysis of the transfer of a group, e.g. a methyl group, glycosyl group, acyl group, phosphorus-containing, or other groups, from one compound (generally regarded as the donor) to another compound (generally regarded as the acceptor). Transferase is the systematic name for any enzyme of EC class 2. | 10 |
| GO:0005515 | Binding to a protein. | 9 |
| GO:0016491 | Catalysis of an oxidation-reduction (redox) reaction, a reversible chemical reaction in which the oxidation state of an atom or atoms within a molecule is altered. One substrate acts as a hydrogen or electron donor and becomes oxidized, while the other acts as hydrogen or electron acceptor and becomes reduced. | 9 |
| GO:0042802 | Binding to an identical protein or proteins. | 6 |
| GO:0016772 | Catalysis of the transfer of a phosphorus-containing group from one compound (donor) to another (acceptor). | 5 |
| GO:0140096 | Catalytic activity that acts to modify a protein. | 4 |
| GO:0016746 | Catalysis of the transfer of an acyl group from one compound (donor) to another (acceptor). | 4 |
| GO:0016875 | Catalysis of the joining of two molecules via a carbon-oxygen bond, with the concomitant hydrolysis of the diphosphate bond in ATP or a similar triphosphate. | 4 |
| GO:0016462 | Catalysis of the hydrolysis of a pyrophosphate bond (diphosphate bond) between two phosphate groups. | 4 |
| GO:0004812 | Catalysis of the formation of aminoacyl-tRNA from ATP, amino acid, and tRNA with the release of diphosphate and AMP. | 3 |
| GO:0016874 | Catalysis of the joining of two molecules, or two groups within a single molecule, using the energy from the hydrolysis of ATP, a similar triphosphate, or a pH gradient. | 3 |
| GO:0016887 | Catalysis of the reaction: ATP + H2O = ADP + H+ phosphate. ATP hydrolysis is used in some reactions as an energy source, for example to catalyze a reaction or drive transport against a concentration gradient. | 3 |
| GO:0003677 | Any molecular function by which a gene product interacts selectively and non-covalently with DNA (deoxyribonucleic acid). | 2 |
| GO:0016788 | Catalysis of the hydrolysis of any ester bond. | 2 |
| GO:0008233 | Catalysis of the hydrolysis of a peptide bond. A peptide bond is a covalent bond formed when the carbon atom from the carboxyl group of one amino acid shares electrons with the nitrogen atom from the amino group of a second amino acid. | 2 |
| GO:0016798 | Catalysis of the hydrolysis of any glycosyl bond. | 1 |
| GO:0016817 | Catalysis of the hydrolysis of any acid anhydride. | 1 |

| textbfGO term | **Definition** | **Count** |
|---|---|---|
| GO:0016757 | Catalysis of the transfer of a glycosyl group from one compound (donor) to another (acceptor). | 1 |
| GO:0016747 | Catalysis of the transfer of an acyl group, other than amino-acyl, from one compound (donor) to another (acceptor). | 1 |
| GO:0017111 | Catalysis of the reaction: a ribonucleoside triphosphate + H2O = a ribonucleoside diphosphate + H+ + phosphate. | 1 |
| GO:0016684 | Catalysis of an oxidation-reduction (redox) reaction in which the peroxide group acts as a hydrogen or electron acceptor. | 1 |
| GO:0042578 | Catalysis of the reaction: RPO-R' + H2O = RPOOH + R'H. This reaction is the hydrolysis of any phosphoric ester bond, any ester formed from orthophosphoric acid, O=P(OH)3. | 1 |
| GO:0046790 | Binding to a virion, either by binding to components of the capsid or the viral envelope. | 1 |
| GO:0008168 | Catalysis of the transfer of a methyl group to an acceptor molecule. | 1 |
| GO:0140098 | Catalytic activity that acts to modify RNA. | 1 |
| GO:0004518 | Catalysis of the cleavage of ester linkages within nucleic acids. | 1 |
| GO:1901363 | Binding to heterocyclic compound. | 1 |
